# One-time learning in a biologically-inspired Salience-affected Artificial Neural Network (SANN)

**DOI:** 10.1101/726331

**Authors:** Leendert A Remmelzwaal, George F R Ellis, Jonathan Tapson

## Abstract

In this paper we introduce a novel Salience Affected Artificial Neural Network (SANN) that models the way neuromodulators such as dopamine and noradrenaline affect neural dynamics in the human brain by being distributed diffusely through neocortical regions. This allows one-time learning to take place through strengthening entire patterns of activation at one go. We present a model that accepts a *salience signal*, and returns a *reverse salience signal*. We demonstrate that we can tag an image with salience with only a single training iteration, and that the same image will then produces the highest reverse salience signal during classification. We explore the effects of salience on learning via its effect on the activation functions of each node, as well as on the strength of weights in the network. We demonstrate that a salience signal improves classification accuracy of the specific image that was tagged with salience, as well as all images in the same class, while penalizing images in other classes. Results are validated using 5-fold validation testing on MNIST and Fashion MNIST datasets. This research serves as a proof of concept, and could be the first step towards introducing salience tagging into Deep Learning Networks and robotics.

## 1. Introduction

This paper introduces a new kind of artificial neural network (ANN) architecture, namely a *Salience Affected Artificial Neural Network (SANN)*. The SANN models the effect of neuro-modulators in the cortex: an important feature of the human brain, based on the well established fact that emotions play a key role in brain function, see for example the writings by Antonio Damasio such as *Descarte’s Error* [3] and *The Feeling of What Happens* [4]. This SANN architecture gives powerful additional functionality to SANNs that are not possessed by other ANNs.

### 1.1. One-time salience tagging

Firstly, this SANN architecture allows for *one-time salience tagging of memories* to take place, by affecting entire patterns of activation in one go. A memory trace of a specific event is formed and stored together with a salience tag associated with the event, which could be positive or negative, depending on its nature. When a triggering event takes place, for example seeing a man approach threateningly with a knife in his hand, one experiences the corresponding perceptions (sight, sound, etc) together with a salience (or significance) signal which focuses attention on that event and associated features. That salience signal, experienced as emotional feelings, is an indication that these are important issues for welfare or even survival that must be dealt with right now. The mind puts other issues aside, and decides how to handle this event in a safe way. This is the immediate effect on attention, and so changes cognitive processes by causing attention to have a specific focus (thinking about a knife rather than walking to the bus stop).

One of the ways in which memories are tagged with salience in the cortex is via changes in synaptic weights due to neuromodulators that are spread diffusely from the excitatory system (based in pre-cortical areas) to the neocortex via ascending system (see Fig 1). The one-time learning with this SANN architecture contrasts crucially with the thousands if not millions of repetitions needed to train a neural network to correctly classify objects via back-propagation and similar methods of adjusting neural network weights, as in usual ANNs. For salience to be applied, the SANN first needs to be trained to classify objects, at least with a few epochs of training. Assuming a base layer of classification training to be complete, the salience training is an alternative to additional iterations of back propagation. Salience has a positive impact on both the individual and the class, assuming that a base level of classification training has already taken place. The mechanism whereby this happens will be discussed in depth below.

**Fig 1.**
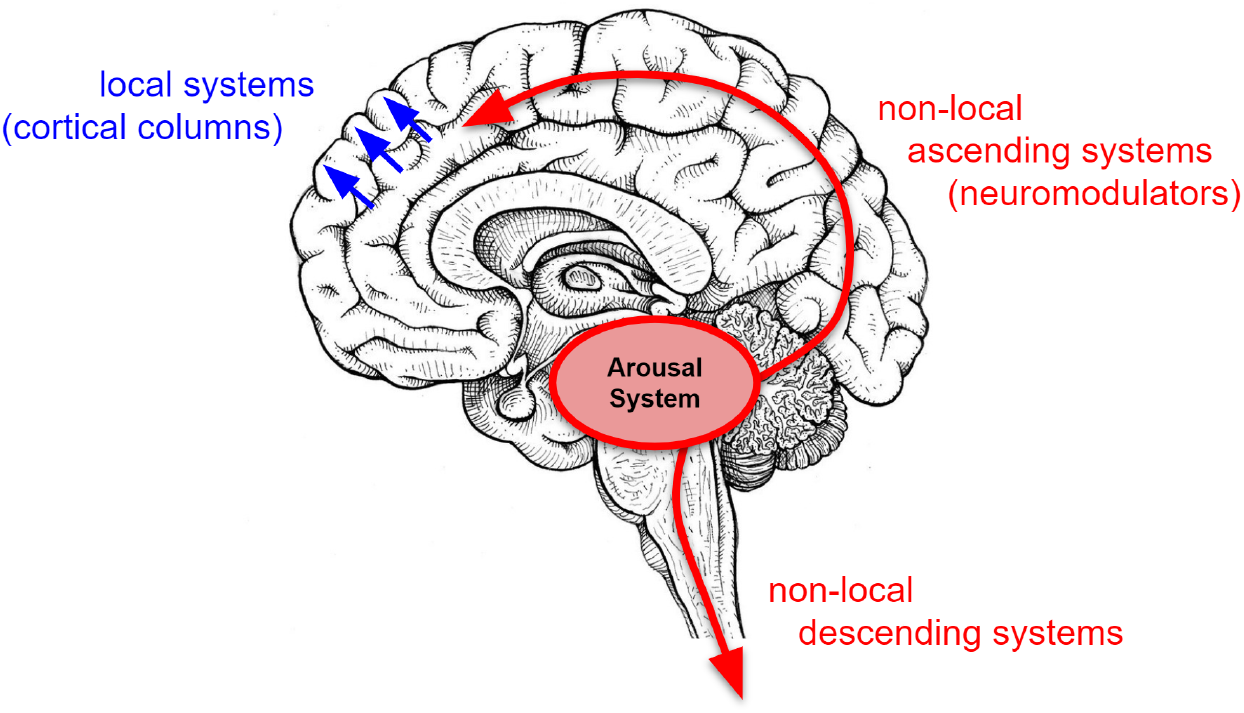
There are two very different kinds of circuits at work in the cortex that we consider here. Firstly there are local systems, such as signals sent between layers in cortical columns. But there are also non-local systems ascending from the arousal system (e.g. the limbic system). The arousal system sends neuromodulators (e.g. dopamine and noradrenaline) into the cortex by means of diffuse projections. These neuromodulators simultaneously affect entire patterns of activated neurons at the time the neuromodulators are delivered, acting on the memory of the object currently being observed.

It is important to note that the one-time learning considered here is not the same as ‘one-shot’ learning [5] [6] which relates to learning categories of entities or events. In contrast ‘one-time’ learning in this paper refers to the tagging of specific individual memory with salience, so that a specific memory is associated with an emotional tag.

### 1.2. Salience retrieved with memories

Secondly, whenever a salience-tagged memory is retrieved, say due to seeing the same person again, the salience tag is retrieved along with the memory, alerting the cortex to its significance and priming it to again act appropriately in response. This reverse salience signal thus gives key guidance as to future actions whenever the memory is reactivated. The SANN architecture allows emotional tags to be attached to sensory inputs, signifying their importance and how to react to them, and recovers these emotional signals when the image is identified an another occasion.

### 1.3. Salience improving learning

Thirdly, the SANN architecture models the impact that diffuse neuromodulators in the cortex have on neurons; affecting the activation functions of neurons, as well strengthening the synaptic weights. This study demonstrates how a salience signal can improve the classification accuracy of the salience-tagged image, as well as improving the classification accuracy of images in the same class. What we also discover is that tagging certain images with salience has a penalizing effect on the classification accuracy of images belonging to other classes.

### 1.4. Scope of this paper

This paper investigates all three aspects noted above. After describing this architecture, the features noted will be demonstrated by a systematic exploration of examples, using the publicly available MNIST data base of handwritten letters so that others can similarly test the model should they wish to do so. It could of course for example be used with a data base of face images instead.

### 1.5. Key features of the SANN

The key feature modelled here is the way neuromodulators such as dopamine and noradrenaline are distributed diffusely through neocortical regions by ‘ascending systems’ originating in nuclei in pre-cortical areas (see Fig 1). These connections contrast with the highly specific synaptic connections between neurons in neocortical columns, which of course also occur and are modelled in standard ANNs. The ascending systems are not connections to specific neocortical neurons: rather they spread neuromodulators to all synapses in specific cortical regions. The neuromodulators then affect an entire pattern of synaptic connections that are active in that region at that time by altering their weights in proportion to the product of the synaptic level of activity and the strength of the neuromodulator released. They leave unaltered weights of all synapses that are inactive. This is the extremely powerful mechanism, described in the book *Neural Darwinism* by Gerald Edelman [7], which strengthens an entire pattern of cortical activation at one go. That is what enables one-time learning to occur. The link to emotions, and so affect, is because these ascending systems are also the physiological basis of the genetically determined primary emotional systems described in detail in the book *Affective Neuroscience* by Jaak Panksepp [8].

### 1.6. The basic underlying assumptions

There are two basic assumptions underlying this paper. Firstly, evolution has fine-tuned human brain structure over millions of years to give astonishing intellectual capacity. It must be possible for designers of ANNs to learn possible highly effective neural network architectures from studying brain structure. This has of course happened in terms of the very existence of ANNs, which are based in modeling the structure of cortical columns. It has not so far happened as regards the structure and function of the ascending systems considered here (an exception is Edelman himself who modelled them, but he left out key aspects as we discuss below). However they have been hardwired into the human brain by evolutionary processes precisely because they perform key functions that have greatly enhanced survival prospects. This architecture should therefore have the capacity to significantly increase performance of any kind of ANN, and so has the potential to play an important role in robotics or AI.

Secondly, while the brain is immensely complex and therefore requires study at all scales of detail in order that we fully understand it, nevertheless it can be claimed that there are basic principles that characterise its overall structure and function, that can be very usefully developed in simple models such as presented here. These models can provide an in-principle proof that the concept works, and so it may be worth incorporating this structure in much more complex models such as massive deep learning or reinforcement learning networks. The testing we do on the simple models presented here suggests that may indeed be the case.

### 1.7. Limitations

This paper does not model either what triggers the salience signal, or its direct effects on cognition and attention. These undoubtedly need to be incorporated in a more complete model of the brain processes of interest, but in order for our investigation to be focused and manageable we concentrate on the effects mentioned above. The relation to perception and attention will be the subject of future papers. We also do not attempt to model spiking neural networks; that again can be the subject of future investigation.

### 1.8. Related Work

There have been a number of attempts to model non-local effects in neural networks and robotics. In this section we review related works.

#### Husbands’ model of gas diffusion

The only other class of ANNs of which we are aware that represent non-local effects are models of the effects of Nitric Oxide (NO), called GasNets [9]. GasNets model the presence of the NO gas in the environment surrounding the neurons, which is capable of non-locally modulating the behaviour of other nodes. Like SANNs, this form of modulation allows a kind of plasticity in the network in which the intrinsic properties of nodes are changing as the network operates [9]. However to the best of our knowledge, GasNets have not been modified to train and test an ANN with specific salience, nor have they been used to demonstrate one-time learning in ANNs. The SANN model differs from Husbands’ GasNets [9] as it models both the way memories associated with emotionally-laden events may be embodied in neural networks, and the way those emotional associations can be recalled when the events are remembered. Furthermore, this research introduces an additional input salience signal during training, and each node produces a nodal reverse salience signal during testing. This research also demonstrates that SANNs are capable of one-time learning by associating strong salience signals with an input combination.

#### Juvina’s model of valuation and arousal

Juvina [10] presents an approach to adding primitive evaluative capabilities to a cognitive architecture in which two sub-symbolic quantities called ‘valuation’ and ‘arousal’ are learned for each declarative memory element, based on usage statistics and the reward it generates. Consequently each memory element can be characterized as positive or negative and having a certain degree of affective intensity. In turn, these characteristics affect the latency and probability of retrieval for that memory element. This is similar to what we do, but using a very different cognitive architecture. The SANN architecture is based closely on the way the diffuse systems interact with cognitive functioning in the human brain [7] [11], which is an affective process because these systems are those implicated in the primary emotional systems identified by Panksepp [8].

#### Edelman’s Robots

Gerald Edelman created a range of brain-based devices (BBDs) including Darwin VII and Darwin X. [2] [12]. Edelman created ‘Darwin VII’ to capture a holistic picture of its environment, incorporating 3 different senses (vision, auditory, conductance or “taste”), a motor system, and a “value system” (attempting to model an ascending neuromodulatory value or reward system). Edelman then attempted to model long-term episodic memory (behaviour attributed to the hippocampus) in his model ‘Darwin X’. Edelman’s robot ‘Darwin VII’ contained a computational nervous system of 20,000 neuronal units, and 450,000 synaptic connections. After some training ‘Darwin VII’ successfully learned to associate the taste value of the blocks with their visual patterns [2]. The sensory systems interpreted incoming signals (such as conductance of a block) as a ‘value signal”. In Edelman’s model of the value system, the strength of connections between two neural units could change based on the value signal (i.e. a positive value signal would strengthen a connection).

Key difference relative to our paper: Edelman allowed the ‘salience’ signal in ‘Darwin VII’ to directly impact the strength of synaptic connections between neurons. By doing so, Edelman’s model incorporates ‘salience’ as just another input signal (along with the visual, auditory and sensory signals) that requires extensive training. As a result, Edelman’s BBD models required many iterations of training for the salience to be embedded in the model, and associated with sensory input patterns. Due to the design of his model, Edelman’s model was not able to demonstrate one-time learning on an previously trained dataset.

##### 1.8.1. Attention

A similar concept to ‘salience’ is the ‘attention mechanism’; a mechamism used commonly with Neural machine translation (NMT) [13]. Attention mechanisms applied to translation activities can improve translations by selectively focusing on parts of the source sentence during translation [14] [15]. This is usually achieved by swapping out specific layers in the neural network with alternative neural network layers, for example Vaswania replace a convolutional layer with a multi-head/self-attention ‘Transformer’ layer [16]). These attention layers also have hidden representations that scale in size with the size of the source [17].

By comparison, the SANN does not swap out or scale hidden representation layers dynamically. The SANN implements an additional salience variable on top of a standard ANN, without changing the dimensions of the hidden representations. In addition the salience signal applies to every node in the neural network (excluding the input layer), rather than applying only to one layer in the neural network. Furthermore, the SANN is able to absorb the salience associated with a memory in a single iteration of one-time learning, and then produce a similar reverse salience signal when the same (or similar) event occurs in the future. While attention networks are similar in some ways, SANN architecture is fundamentally different.

#### The SANN model

In contrast to other related work, in this paper we demonstrate how the ‘salience’ signal from the arousal system (which Edelman called the ‘value’ system) can be modelled as an additional dimension to the neural network model. We demonstrate how a salience signal can affect a specific neural activation patterns (e.g. a memories), tagging it with a salience signal (e.g. an emotional response) during one-time learning. It is important to note that this research is not the same as some “emotional robots” that are constructed so as to physically simulate emotional expressions. We rather simulate the way affect alters cortical processing. One could additionally add the aspect of emotional expression simulation if desired, but this falls outside of the scope of this research.

### 1.9. Structure of this paper

In section 2 we describe how we embed a *salience signal* and a *reverse salience signal* in a neural network. In section 3 we describe the experimental design, and in Section 4 we present the results of the simulations we ran and the observations made. We then draw conclusions in section 5, and make suggestions for future work in section 6.

## 2. Modelling salience

In this section we discuss the implementation of the salience signal (*S*) and reverse salience signal (*R*) in a neural network. The overall architecture is shown in Fig 2 and described in detail in this section.

**Fig 2.**
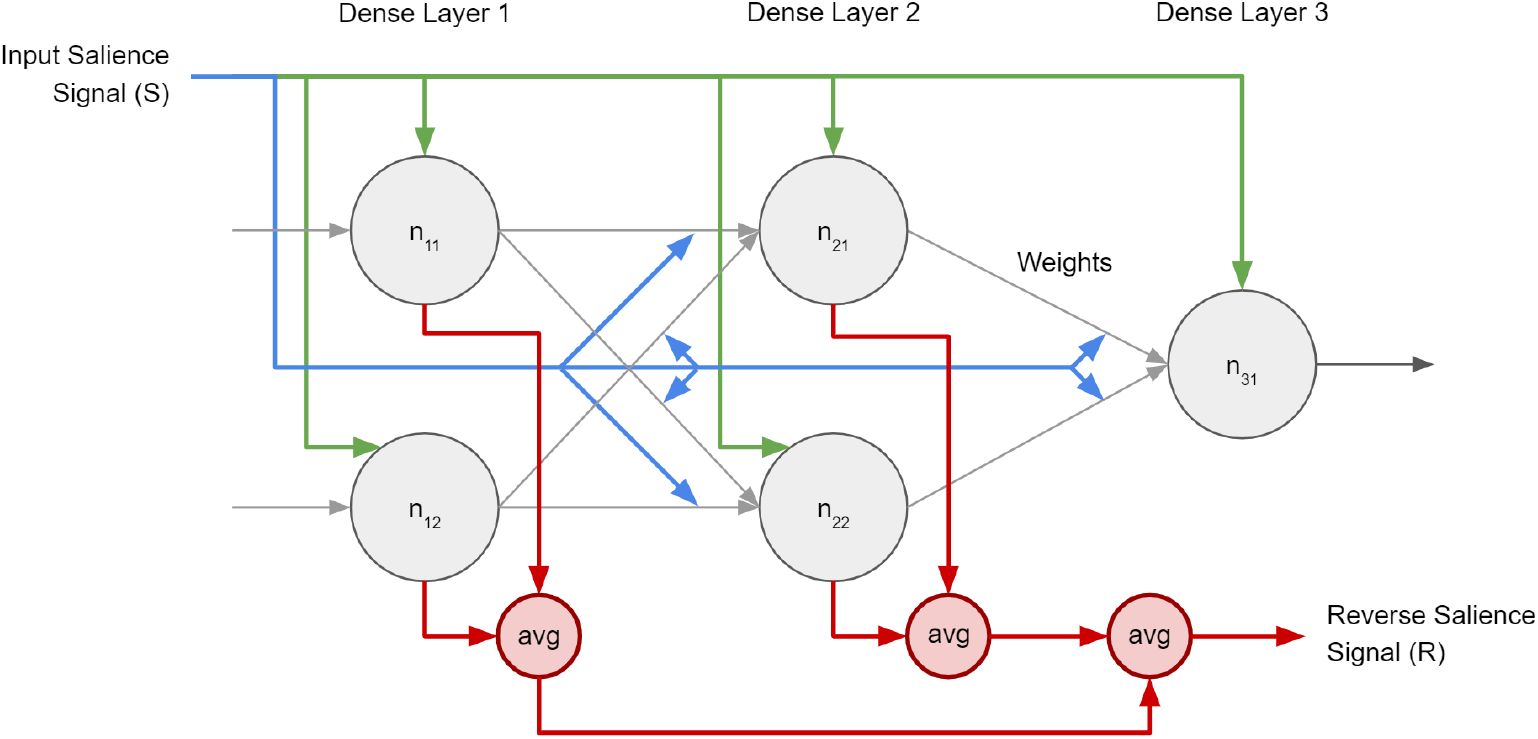
The SANN is an ANN with an additional signal dimension added: the salience signal. The salience signal (S) affects all nodes (green arrows) and weights (blue arrows) in the network simultaneously during training. Each node in the network then produces a reverse salience signal (R) during testing. Reverse salience signals (red arrows) are averaged for each layer, and all Dense Layers are averaged to calculate the global reverse salience signal.

### 2.1. Salience signal (S)

The salience signal (*S*) is a global signal that applies uniformly to each node in the neural network during salience training, in the same way that the presence of a neuromodulator (such as dopamine) affects a group of neurons similarly. To implement a salience signal we created a new salience variable (*S*) for each node in each densely connected layer in the network. During a salience training iteration the network was presented with a global salience signal (*S*), which affected each node (as seen in Fig 2).

The salience value of each node (*S*) is a value between −1 and 1. The change of salience value of each node during salience training is proportional to the activation levels of the post-synaptic node (*A*) and the global training salience value (*S*), as shown in Equation 1.

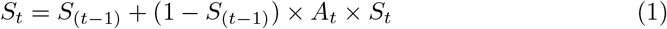

### 2.2. Reverse salience signal (R)

In addition to a global salience signal (*S*), we also create a global reverse salience signal (*R*). While the global salience signal (*S*) affects nodes during salience training, the reverse salience signal (*R*) is produced by the neural network at the time of classification, alongside the classification output. The reverse salience signal for each node (*R*_node_) was calculated as the product of the activation (*A*) of the node, and the salience value (*S*) of the node (see Equation 2). The reverse salience for each densely-connected layer was then calculated as the average reverse salience in the layer (*R*_layer_), seen in Equation 3). The reverse salience for the network (*R*_layer_) was calculated as the average of the reverse salience from each layer (*R*_layer_), seen in Equation 4.

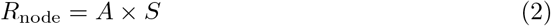

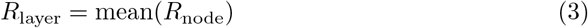

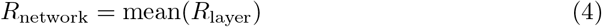

This was done so that each layer received an equal contribution to the total reverse salience of the network; averaging all the nodes in the network would have given less weight to layers with fewer nodes (as seen in Fig 2). Initially the salience signal was designed as an independent signal. In the next section we explore the impact of the salience signal on learning.

### 2.3. Impact of Salience on Learning

Biologically speaking, adding salience to an event or memory has a lasting impact on the quality of that memory, in addition to producing a reverse salience signal upon recalling the event. For example, if someone is bitten by a dog, then a heightened fear of dogs in general is expected (*class*), and even higher fear is expected when they see the specific dog that bit them (*specific*). However, in addition we would also expect that having had an emotional event associated with a dog (i.e. being bitten) would improve that person’s ability to classify dogs in general, but especially the specific dog that bit them.

In this paper we aim to demonstrate that the addition of salience can improve the accuracy of classifying a *specific* input, as well as similar elements in the same *class*. For example, if someone is bitten by a dog, then a heightened fear of dogs in general is expected (*class*), as well as an even higher fear of the specific dog that bit them (*specific*). This distinction is shown in Fig 3 below.

**Fig 3.**
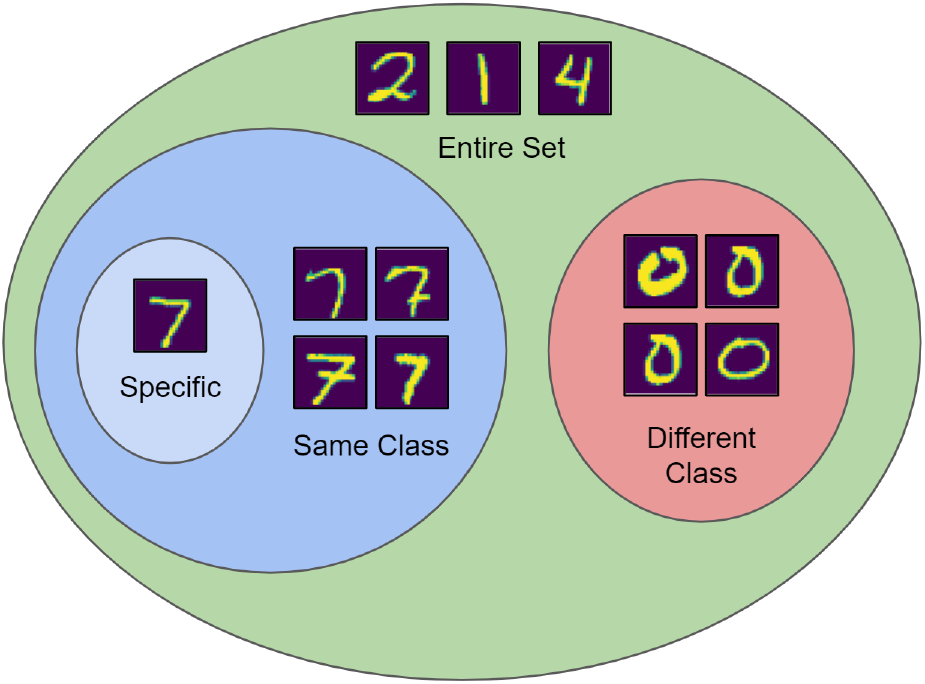
We identify that an input can belong to one of 4 classes within a dataset: (1) specific image that was tagged with salience, (2) same classification class as the tagged image, (3) a different classification classes to the tagged image, and (4) the group of all images in the entire dataset.

In this section we investigate the effect of the the salience value (*S*) on (a) the activation functions of each node, as well as (b) the strength of the weights in the network. We will review these effects in this section.

#### 2.3.1. Impact on Activation Functions

To closely model the behaviour of neuromodulators on biological neurons, we explored the impact of salience on the activation functions of each node. We chose to use the sigmoid activation function (described in Equation 5), because it was better suited than other activation functions (e.g. maxout functions [18], ReLU functions [19], PLUs [20]) to the SANN model for two reasons. Firstly, because sigmoidal activation functions are more biologically realistic: most biological systems saturate at some level of stimulation (where activation functions like ReLU do not). And secondly, sigmoidal functions allow for bipolar activations, whereas functions like ReLU are effectively monopolar and hence not useful for a reverse salience signal with both polarities of activation.

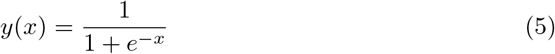

We chose to investigate three possible modifications to the activation function, namely (a) a horizontal offset, (b) a change in the gradient, and (c) a change in the amplitude. These variations have been visualized in Fig 4.

**Fig 4.**
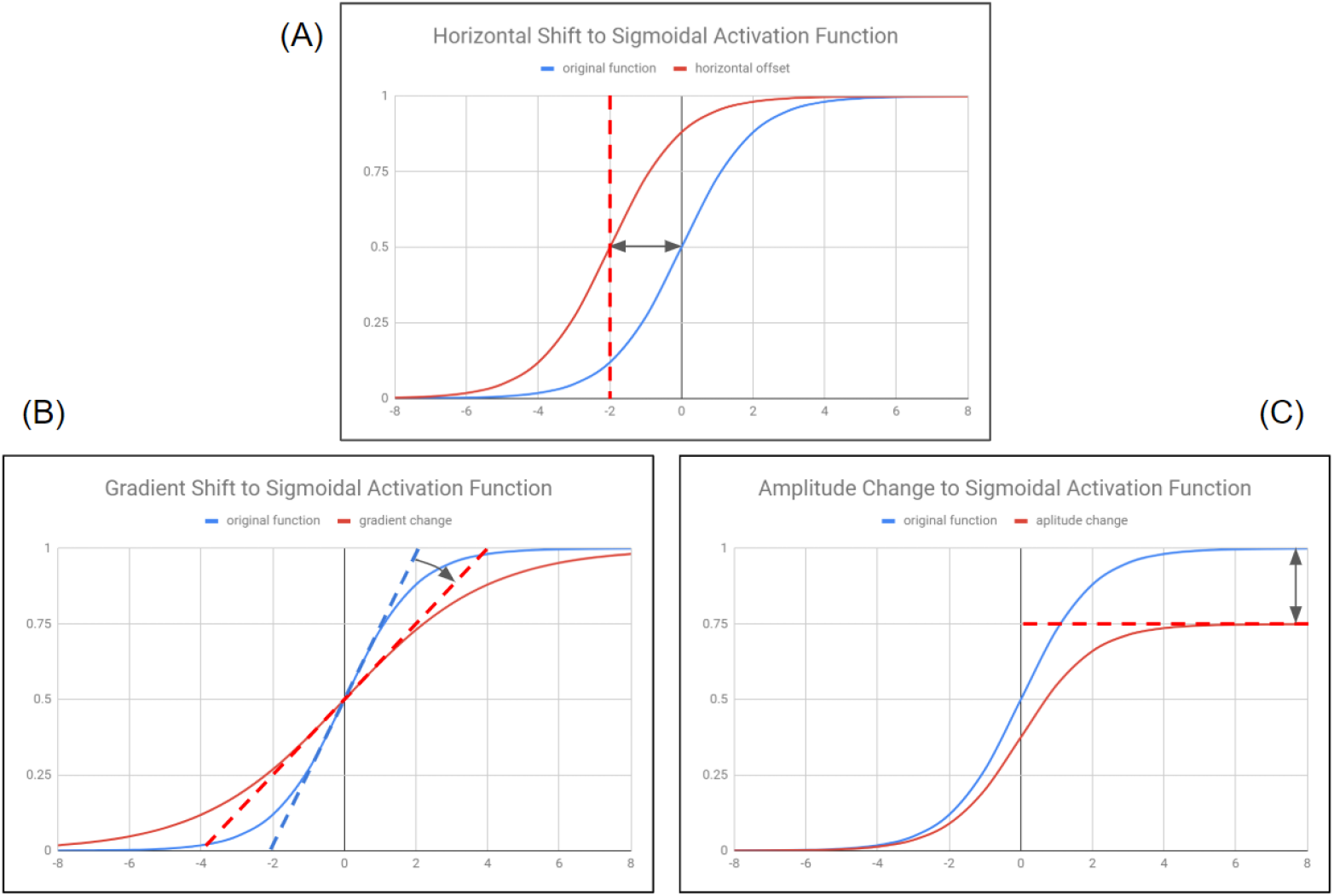
Three changes to the activation function were explored: (A) Change in the horizontal offset of the activation function: A positive salience signal would shift the activation function to the left, resulting in a higher output from the activation the next time. (B) Change in the gradient of the activation function: A positive salience signal would reduce the gradient of the activation function, resulting in a higher output from the activation the next time. (C) Change in the amplitude of the activation function: A positive salience signal would increase the amplitude of the activation function, resulting in a higher output from the activation the next time.

##### Horizontal Offset

The first of three effects was to introduce a horizontal shift of the activation function along the x-axis (see Fig 4A). A positive salience signal would shift the activation function along the x-axis, resulting in either a higher or lower output from the activation the next time. We explored the effects of a positive shift (shown in Equation 6) as well as a negative shift (shown in Equation 7).

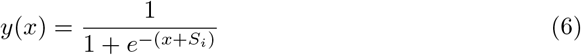

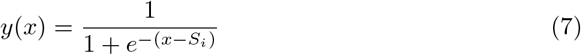

##### Gradient Change

The second of three effects was to introduce a change in the gradient of the activation function (see Fig 4B). A positive salience signal would reduce the gradient of the activation function, resulting in a higher output from the activation the next time. The activation function incorporated the gradient change as shown in Equation 8.

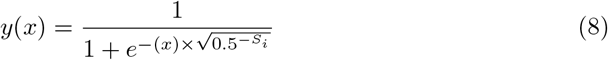

##### Amplitude Change

The third of three effects was to introduce a change in the amplitude of the activation function (see Fig 4C). A positive salience signal would increase the amplitude of the activation function, resulting in a higher output from the activation the next time. The activation function incorporated the amplitude change as shown in Equation 9.

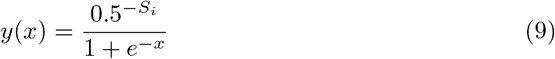

#### 2.3.2. Impact on the Strength of Weights

In addition to producing a reverse salience signal (*R*), the global salience signal (*S*) also has the impact of altering the weights of the connection between all nodes (*W*) in the network simultaneously, by an amount proportional to the current weight of the connection (*W*_*i*_), the strength of the input salience signal (*S*_*i*_), and the activation level of the post-synatic node (*A*_*j*_).

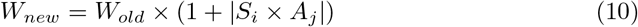

Here *α* is a constant which might differ from neuron type to neuron type, or with the kind or neuromodulator modelled. Note that because the updating equation for each synapse weight depends on the current state of activation (*A*_*i*_) of that synapse, just as in the case of Neural Darwinism [7], it is the entire current pattern of activation that gets reinforced by the salience signal.

## 3. Experimental Design

In this section we discuss the experimental design, including the neural network architecture, training parameters, and datasets.

### 3.1. Neural Network Design

To evaluate the implementation of salience we tested it on a fully-connected 3-layer neural network (ANN). Architectures such as LeNet architecture [21], AlexNet [22], VGGNet [23], GoogLeNet [24] and ResNet [25] [26] were omitted from this study because they utilize a convolutional layer, which is out of scope of this research.

### 3.2. Software implementation

We test our hypothesis using a 3-layer fully-connected artificial neural network (ANN). The dimensions of the ANN are [256-256-10]. The neural network framework was adapted from an open-source pure python implementation of a Neural Network [27]. The activation functions of each node was sigmoidal; variations of the activation function (e.g. ELU, ReLU, Leaky ReLU) could be tested in further research.

### 3.3. Dataset

In this paper we chose to validate our results using 2 datasets (MNIST [28] and Fashion MNIST [29]). The images in the datasets were reshaped to 16 x 16px. Other datasets such as CIFAR10 [30] and GTSRB road sign dataset [31] were considered, but these datasets perform best with a convolutional neural network; adding convolutional layers falls outside of the scope of this research.

### 3.4. K-Fold Validation Testing

The training was limited to the first 1000 images in each dataset. We implemented 5-fold validation testing, so that each training dataset contained 800 images, and each testing dataset contained 200 images.

### 3.5. Fixed Parameters

The fixed parameters for the neural network models include:

- Layers: 3
- Layer Dimensions: 256 (input), 256 (hidden), 10 (output)
- Supervised learning algorithm: Back propagation
- Shuffling the samples: True
- The activation function: Sigmoidal
- Initializing the Weights: Random
- Learning Rate: 0.1
- K-Folds: 5
- Random seed: 0 (to ensure that results are reproducible)

## 4. Experiments and Results

In this section we discuss the various experiments we conducted, namely:

i. the baseline accuracy test
ii. the addition of salience
iii. the calculation of reverse salience
iv. the effect of negative salience
v. the effect of mixed salience training
vi. the effect of reverse salience on the activation functions and weights of each node

### 4.1. Baseline performance

The neural network was trained for 20 epochs, and the baseline performance results for the MNIST and Fashion MNIST datasets are visualized in Fig 6, and the final accuracy metrics are shown in Table 1.

**Table 1.** Baseline classification accuracy after 10 epochs of classification training across MNIST and Fashion MNIST datasets. Mean and Standard deviation calculated using 5-fold validation accuracy.

| Classification accuracy after classification training only |  |  |
| --- | --- | --- |
| Dataset | MNIST | Fashion MNIST |
| Training Iterations | 10 epochs | 10 epochs |
| Baseline | 83.00% ± 3.74% | 70.90% ± 3.18% |

**Fig 5.**
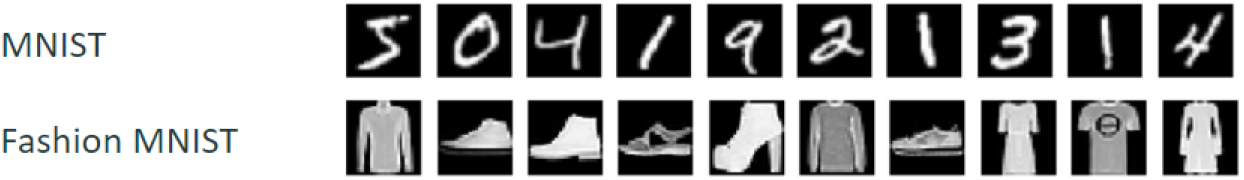
Examples of 10 images (16 × 16px) from MNIST and Fashion MNIST datasets.

**Fig 6.**
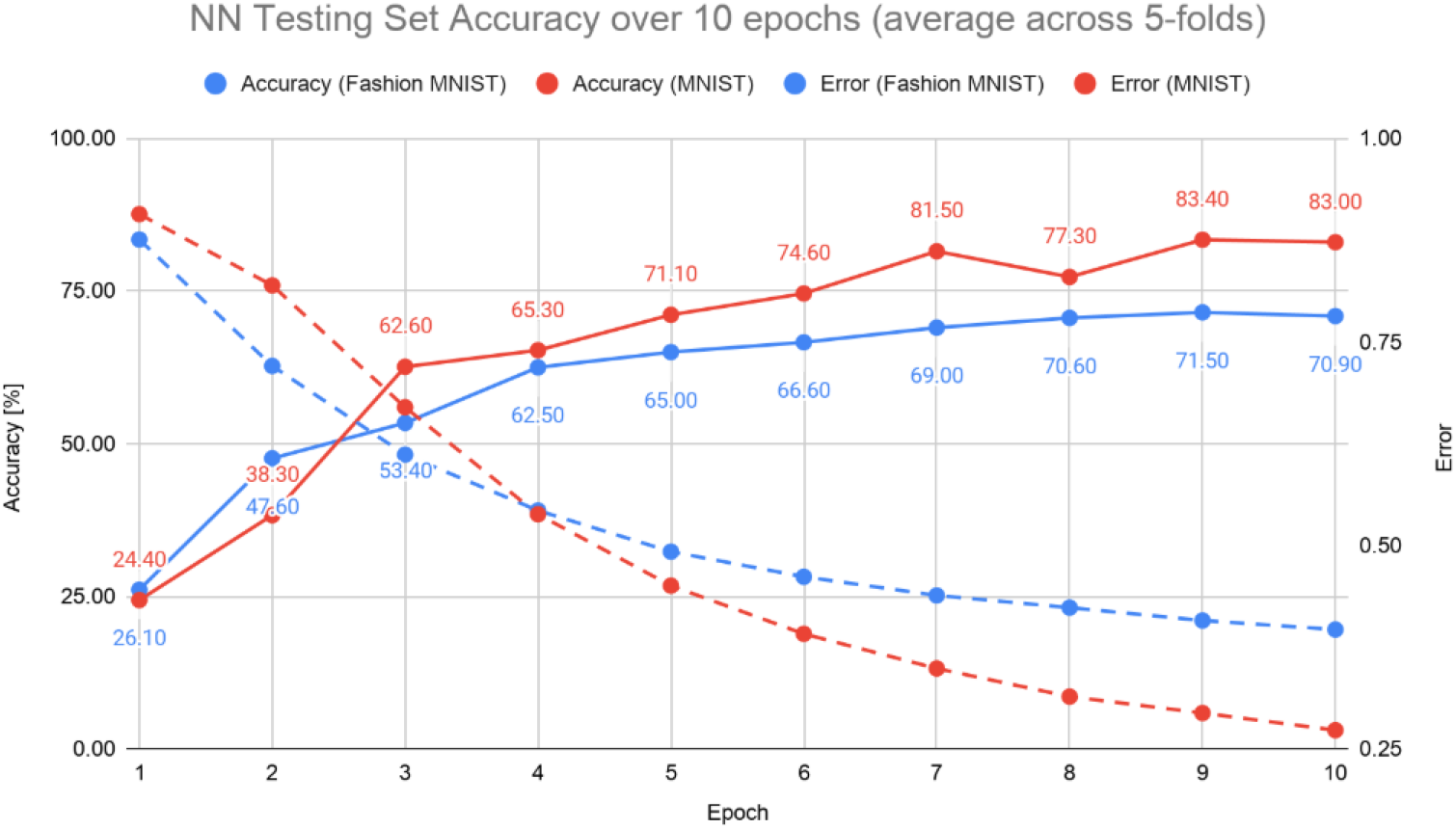
Baseline Accuracy of the MNIST and Fashion MNIST datasets over 10 epochs (average accuracy across 5-folds). Solid lines represent accuracy, and dotted lines represent error.

### 4.2. Salience

We initially added the salience value to each node in the network as an *independent* variable. During training we tagged a single image with positive salience, and ran a single iteration of salience training across the network. The salience value of each node was updated as described in Section 2.1. After a single iteration of salience training, each node in the network (excluding the input layer) was tagged with salience. A visualization of this is shown in Fig 7.

**Fig 7.**
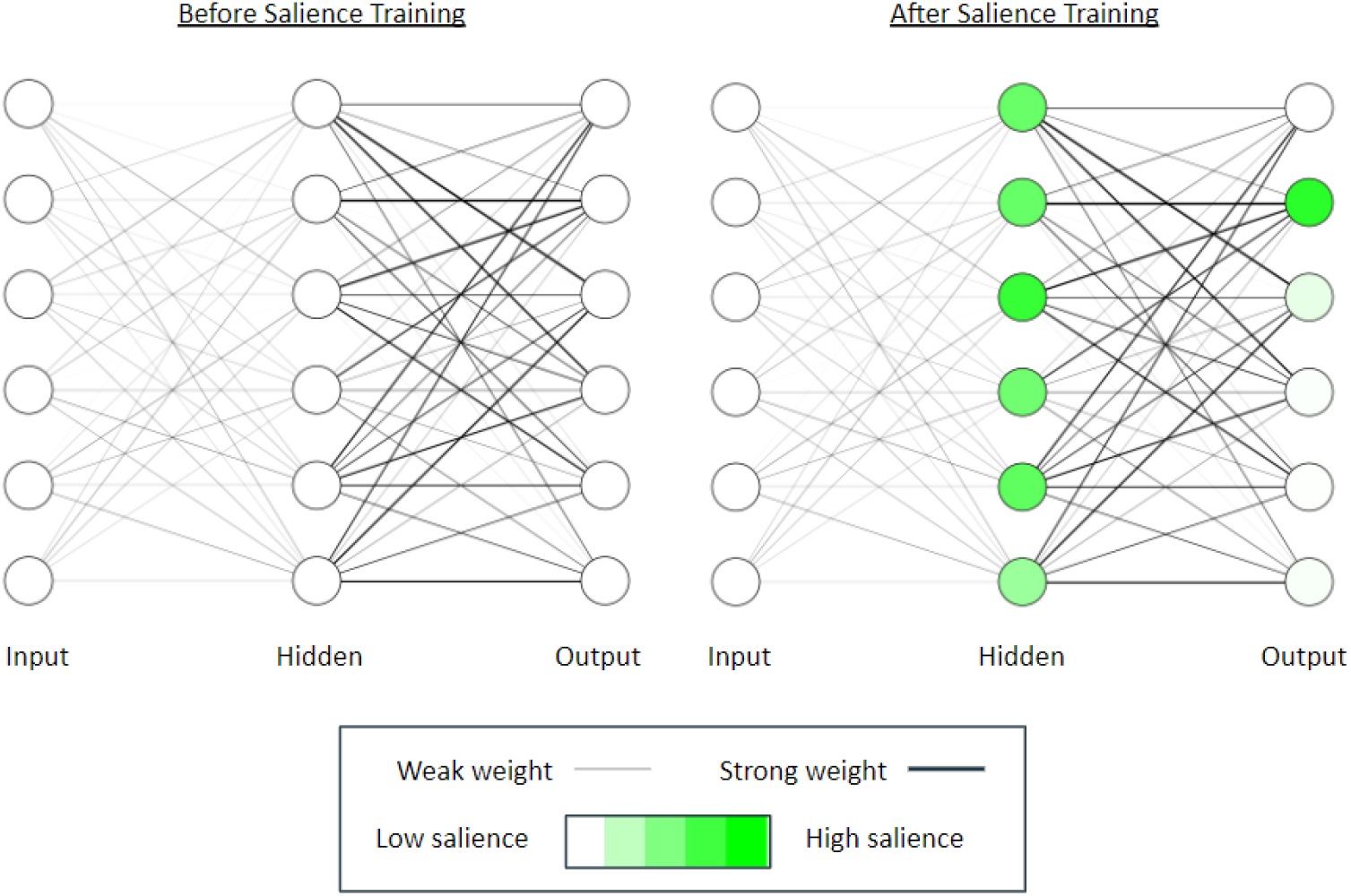
A visualization of the salience values of each nodes of the neural network, before and after one-time positive salience training. The dimensions of the ANN are [256-256-10] but this visualization is limited to 10 layers per layer.

### 4.3. Reverse Salience

Reverse salience was then calculated as the average reverse salience of each node in the output layer of the neural network, as described in Section 2.2. After a single iteration of salience training, the reverse salience was averaged across all 800 elements in the k-fold training set. We observed reverse salience for 4 classes of images within the dataset: (1) specific image that was tagged with salience, (2) the average for all images in the same classification class to the tagged image, (3) the average for all images in the different classification classes to the tagged image, and (4) the average for all images in the entire dataset, as shown in Fig 3. Results are shown in Table 2 and illustrated in Fig 8.

**Table 2.** Reverse salience values produced for both the MNIST and Fashion MNIST datasets after a single iteration of salience training. Mean and Standard deviation calculated using 5-fold validation accuracy.

| Reverse saliency signal after one-time saliency training |  |  |
| --- | --- | --- |
| Dataset | MNIST | Fashion MNIST |
| Classification Training | 10 epochs | 10 epochs |
| Saliency Training Iterations | one-time | one-time |
| Reverse Saliency (Specific Image) | $0.07616 \pm 0.02405$ | $0.07037 \pm 0.02871$ |
| Reverse Saliency (Same Class) | $0.07369 \pm 0.00989$ | $0.06766 \pm 0.01917$ |
| Reverse Saliency (Different Class) | $0.00554 \pm 0.00446$ | $0.00574 \pm 0.00486$ |
| Reverse Saliency (Entire Set) | $0.01283 \pm 0.00422$ | $0.01327 \pm 0.00487$ |

**Fig 8.**
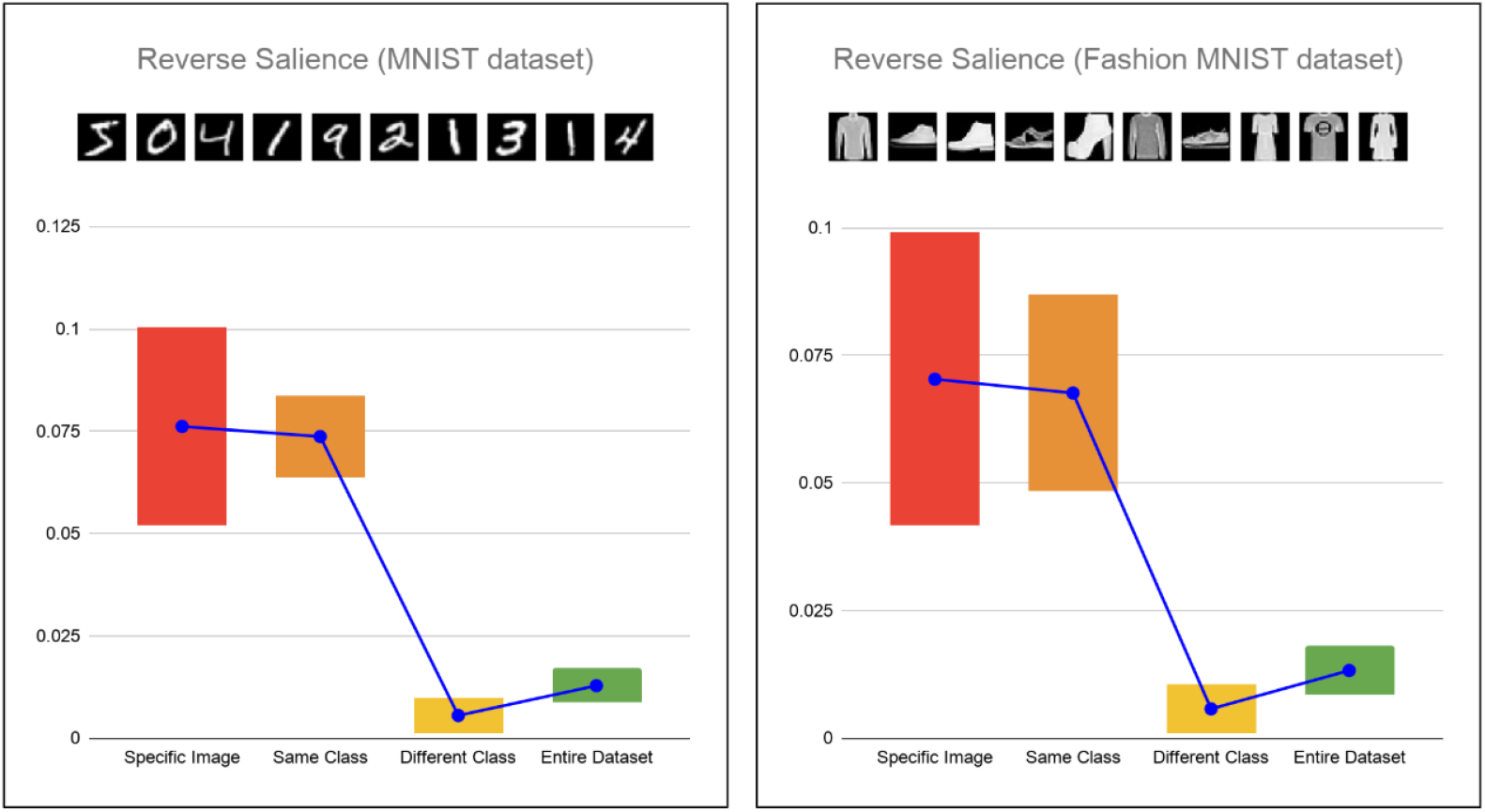
Reverse salience values produced for both the MNIST and Fashion MNIST datasets after a single iteration of salience training. Simulations were run across all 800 elements in the dataset and results were averaged. Results are grouped into 4 classes of images within the dataset: (1) specific image that was tagged with salience, (2) the average for all images in the same classification class to the tagged image, (3) the average for all images in the different classification classes to the tagged image, and (4) the average for all images in the entire dataset. The blue line represents the average, while the bars represent the average +−1 standard deviation. Mean and Standard deviation calculated using 5-fold validation accuracy. These results show that Specific images tagged with salience during training have the highest average, followed by elements in the similar class. Elements in the different class produced the lowest reverse salience, even lower than the average for the entire dataset.

We observed that the specific images tagged with salience had the highest reverse salience, followed by images in a similar class, followed by images in the entire dataset.

The average reverse salience for images in a different class dropped below the average for the entire set.

### 4.4. Negative Salience Training

In this experiment we explore a change in sign of the salience signal. A negative salience signal introduces a second dimension of salience. A positive salience signal could represent a positive emotional tag, while a negative salience signal could represent a negative emotional tag associated with a specific image or experience.

To explore the impact of the *sign* of the salience signal, we trained the neural networks with a salience value of −1, instead of +1. The architecture was designed such that a negative salience signal would be treated symmetrically opposite in sign, and the results showed exactly this: reverse salience signal (*R*) was identical in *magnitude* to positive salience training, but opposite in *sign*.

### 4.5. Mixed Salience Training

Having conducted experiments with positive and negative salience, we then conducted an experiment to combine these two effects. We performed one-time salience training on one image with *positive* salience, and then one-time salience on another image (different class) with *negative* salience. We see from Fig 9 that two (2) separate salience signals can be successfully encoded in the same neural network. This experiment demonstrates that the SANN can accept multiple layers of salience training, even if the salience signals are different in sign.

**Fig 9.**
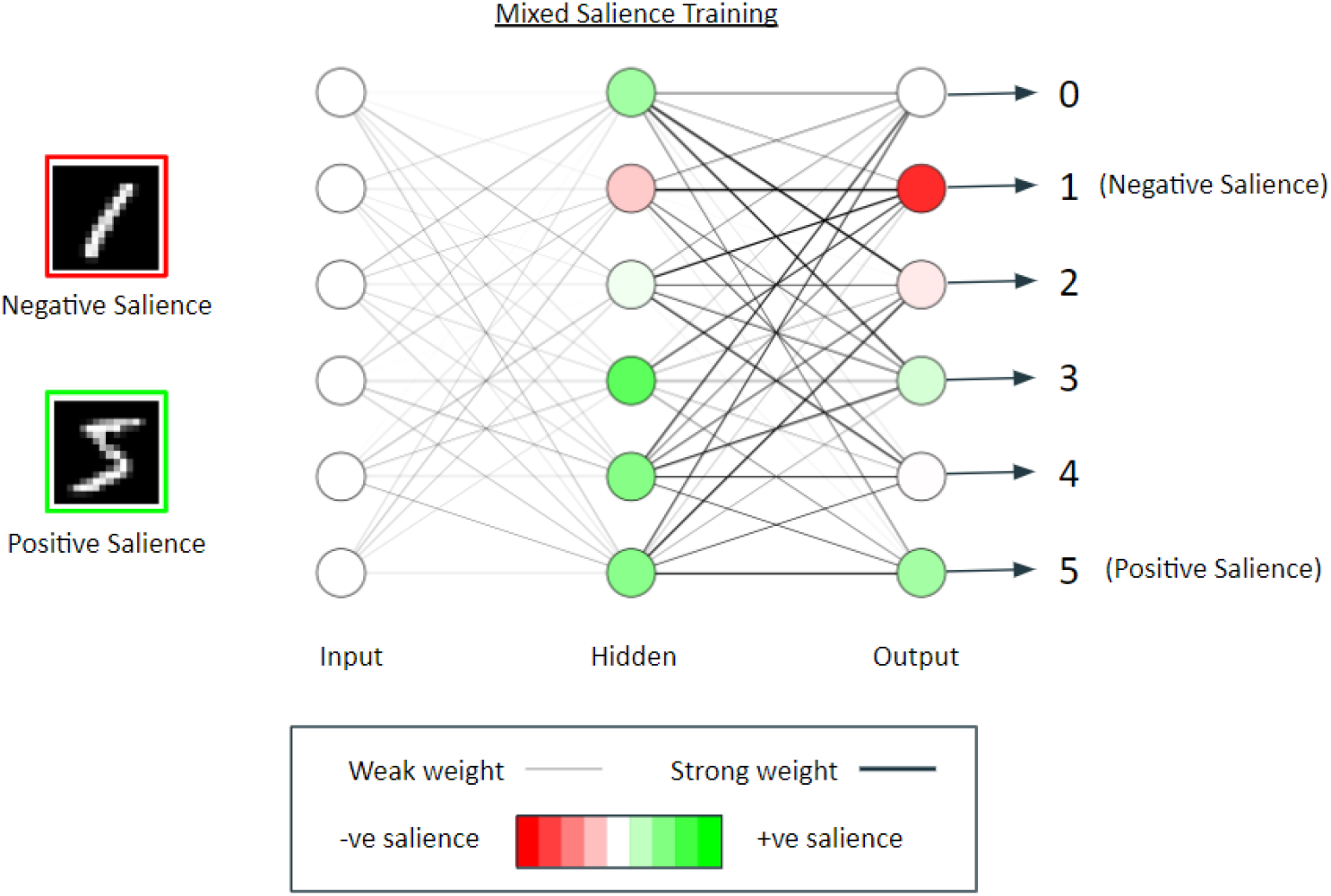
A visualization of the salience values of each nodes of the neural network after mixed salience training: a one-time of positive salience associated with an image in digit class “5”, followed by a one-time of negative salience associated with an image in digit class “1”. The dimensions of the ANN are [256-256-10] but this visualization is limited to 10 layers per layer. We see in this figure that 2 separate salience signals can be successfully encoded in the same neural network.

We suggest that in future research that more than two salience signals are tested.

### 4.6. Impact of salience on learning

In this section we explore the impact of one-time salience training on the accuracy of the neural network. As discussed in the introduction, ‘one-time’ learning in this paper relates to the learning of specific individual entities, rather than the ‘one-shot’ learning of categories of entities or events. We specifically explore the effect of salience on the activation function (gradient change, horizontal shift and maximum value change), as well as the weights in the neural network. We expect the salience to have a similar effect that a neuromodulator in the cortex would have on a specific memory: a improved classification accuracy of the salience-tagged image, as well as all images in the same class as the salience-tagged image.

After the initial classification training had been completed, the baseline classification accuracy was recorded. One-time training was then applied in turn to each image in training set, and reloading the neural network model each time, and the results averaged. Variations were applied to all nodes with salience associated: all nodes in the network except the input nodes. The results from this experiment are shown in Table 3, and an example of weights strengthening is illustrated in Fig 10.

**Table 3.** Classification accuracy of the SANN after one-time salience training across MNIST and Fashion MNIST datasets. Mean and Standard deviation calculated using 5-fold validation accuracy. Bold values indicate an increase in accuracy relative to the baseline score. Allowing the salience signal to impact the Gradient, Horizontal Offset (negative) and the pre-synaptic weights had an overall positive effect on the overall accuracy of the SANN.

| NN Accuracy after one-time saliency training |  |  |
| --- | --- | --- |
| Dataset | MNIST | Fashion MNIST |
| Classification Training | 10 epochs | 10 epochs |
| Baseline | 83.00% ± 3.74% | 70.90% ± 3.18% |
| Saliency Training Iterations | one-time | one-time |
| **Gradient** | **83.22% ± 3.51%** | **72.33% ± 4.10%** |
| Horizontal Offset (positive) | 79.59% ± 4.40% | 62.61% ± 5.74% |
| **Horizontal Offset (negative)** | **83.23% ± 3.34%** | **71.77% ± 4.31%** |
| Amplitude | 82.90% ± 3.60% | 70.88% ± 3.18% |
| **Pre-synaptic weights** | **83.24% ± 3.39%** | **71.52% ± 3.84%** |

**Fig 10.**
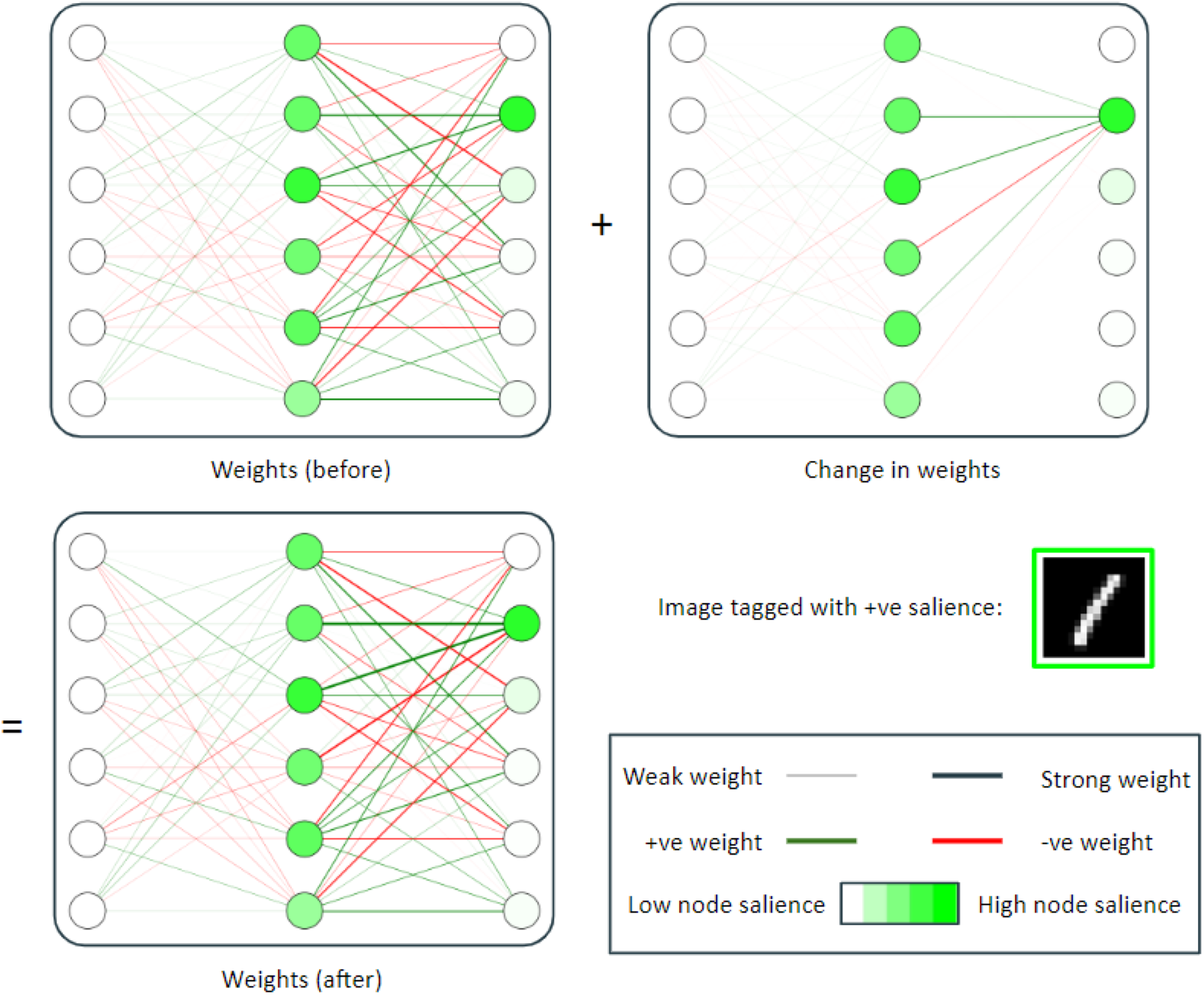
A visualization of the effects of salience strengthening the weights of a specific pattern of nodes relative to the activation of the post-synaptic node. In this example the digit shown (classified as class “5”) was tagged with salience. After forward propagation, the pre-synaptic weights were strengthened proportional to the node activation. The strengthening of the weights in this way has a positive impact on overall accuracy of the network (see Table 3).

### 4.7. Combining impact activation and weights

Finally, we investigated combining the effects of salience on (1) activation function and (2) strengthening the weights. We specifically chose the MNIST dataset, and the change of activation gradient for this simulation. We varied the impact factor on both the activation function and weights, and recorded the overall accuracy of the neural network in the form of a heat map (see Fig 11). From these results we see that there is an optimal band of combined effects that optimizes the overall accuracy. This band ranges from using only the effect on weights (with a factor of 1.2) to only changing the gradient of the activation function (with a factor of 3). However, a combination of these two (weight factor of 0.4 and a gradient factor of 1.5) yields an optimal performance for the entire network.

**Fig 11.**
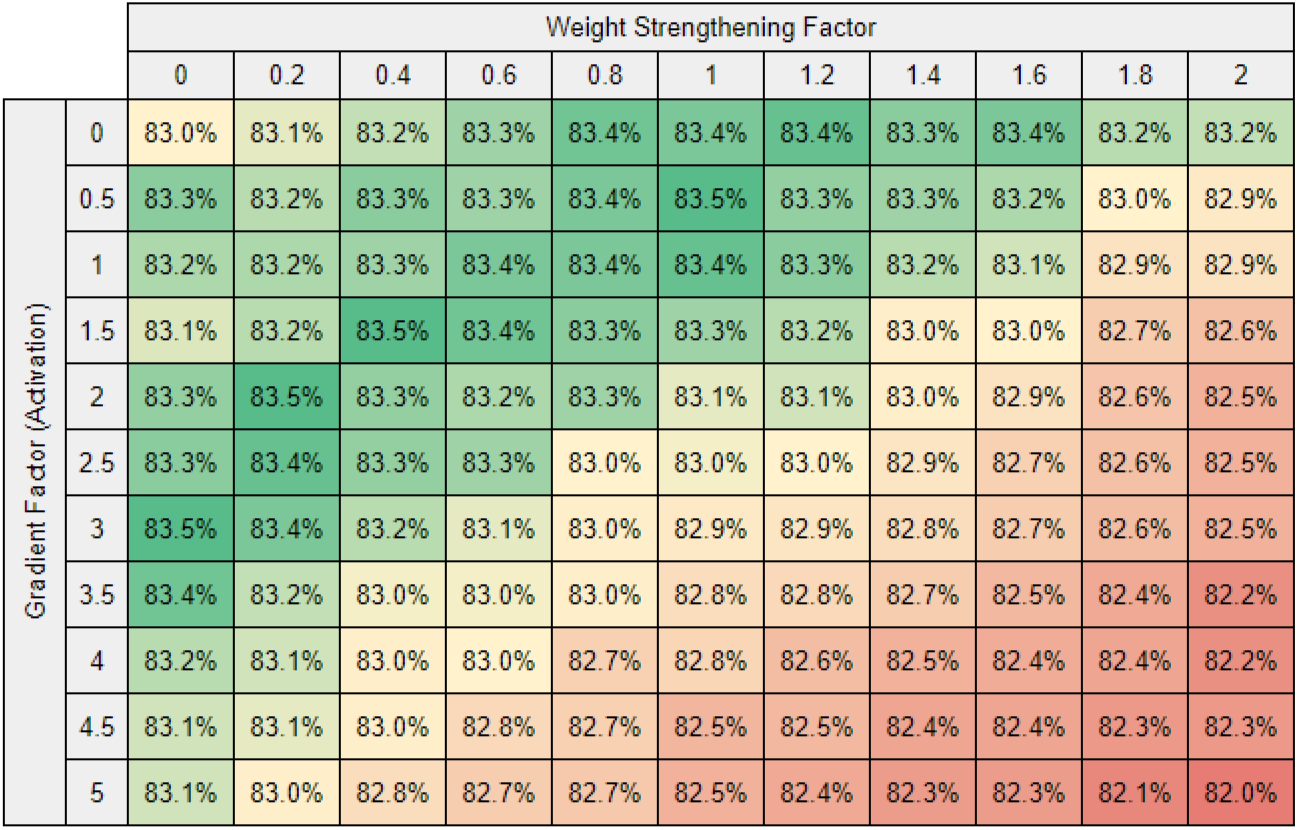
A heat map visualization showing the overall network accuracy as a function of (1) salience changing the gradient of the activation function and (2) salience strengthening the weights. This simulation is limited to the MNIST dataset. Mean and Standard deviation calculated using 5-fold validation accuracy. Standard deviation is not shown in this heat map.

## 5. Discussion

In this paper we have introduces a new kind of Artificial Neural Network architecture, namely a *Salience Affected Artificial Neural Network (SANN)*. The SANN accepts a global salience signal during training, which affects the specific pattern of activated nodes during a single iteration of salience training. This model architecture is inspired by the effects of diffuse projections of neuromodulators in the cortex.

We have demonstrated three key components of one-time salience training in a neural network. Firstly, we have demonstrated how a specific memory can be tagged with *salience* during only one iteration of salience training. Secondly, we have demonstrated that a salience tag can be retrieved along with the memory; a *reverse salience signal*. Thirdly we have demonstrated that salience can improve the classification accuracy of the *salience-tagged image* as well as images in the *same* class, but it also has a penalizing effect on the classification accuracy of images belonging to *other* classes.

To give an example, say we train a neural network to classify images into 3 types of animal classes: cats, dogs, and birds. Furthermore, some of the image are of your favourite cat Cleo. Once the network has been trained to correctly classify images into these 3 classes of animals, you associate a positive salience with your favourite cat Cleo. Associating salience this way in a SANN has 3 distinct impacts: (1) Firstly, the specific memory of your cat Cleo has a heightened reverse salience. (2) Secondly, the entire class of cats also receives a heightened reverse salience signal. (3) Thirdly, all other classes of objects (in this example dogs and birds) experience a relative depression of reverse salience. In other words, associating a positive salience with your cat Cleo has a positive salience impact on all memories of your cat Cleo as well as all other cats, but a relatively reduces the emotion associated with all other animals. It is also important to note that these results show that that there is in fact a ‘penalty’ for learning with salience; salience can have a *positive* impact on some memories, while having a *negative* impact on others.

This paper is limited to demonstrating a proof of concept only; there are many further areas of suggested research, as well as many possible applications of this research, which we will touch on in the next section.

## Glossary

AI: artificial intelligence
ANN: artificial neural network [1]
BBD: brain-based device [2]
MSE: mean squared error
NN: neural network
ReLU: rectified linear unit
SANN: salience-affected neural network

## 6. Future Work

We recommend that in future work the timing of salience training is explored, asking: what effect would there be if salience training took place before or during classification training? The intensity of the salience factor would also be interesting to explore, as would the impact of repeated salience training on the same image, or on a selection of images. Future research may wish to incorporate datasets such as the NIST19 [32], or the Chinese Handwriting dataset [33], and facial datasets such as SCFace [34]. It would be interesting to explore whether the same effect would be observed for other training meta-parameters values (e.g. learning steps, batch sizes, number of layers, etc). Extending this research to other deep neural networks (e.g. recurrent neural networks, convolutional neural networks) is also suggested as future research. These suggestions all fall outside of the scope of work of this initial paper, as this paper is limited to demonstrating a proof of concept only.

## 7. Supporting Material

The source code as well as records of the tests conducted in this paper are publicly available online [36]. For additional information, please contact the corresponding author.

## 8. Acknowledgements

A preliminary version of this work was the subject of an MSc thesis of Remmelzwaal, supervised by Tapson and Ellis. A pre-print was released in 2010 [35]. The present version is so improved and updated that it is essentially a new paper, in particular because, apart from a greatly improved presentation of the logic of the project and its relation to brain structure and function, it has added the dynamics of a shift of weight in proportion to the salience signal and two further effects on the activation function, as well as extensive testing of how this works out in practice.

We are grateful to Mark Solms, Amit Mishra and Jonathan Shock for very helpful comments, and Bruce Bassett for a useful remark.

This research did not receive any specific grant from funding agencies in the public, commercial, or not-for-profit sectors. We declare no conflicting interests.

